## Supporting Information for "Stimulation-induced cytokine polyfunctionality as a dynamic concept"

### Author list and affiliations

**Kevin Portmann[1], Aline Linder[1] and Klaus Eyer[1*]**

[1] *Laboratory for Functional Immune Repertoire Analysis, Institute of Pharmaceutical Sciences, Department of Chemistry and Applied Biosciences, ETH Zürich, 8093 Zürich.*

### Figures

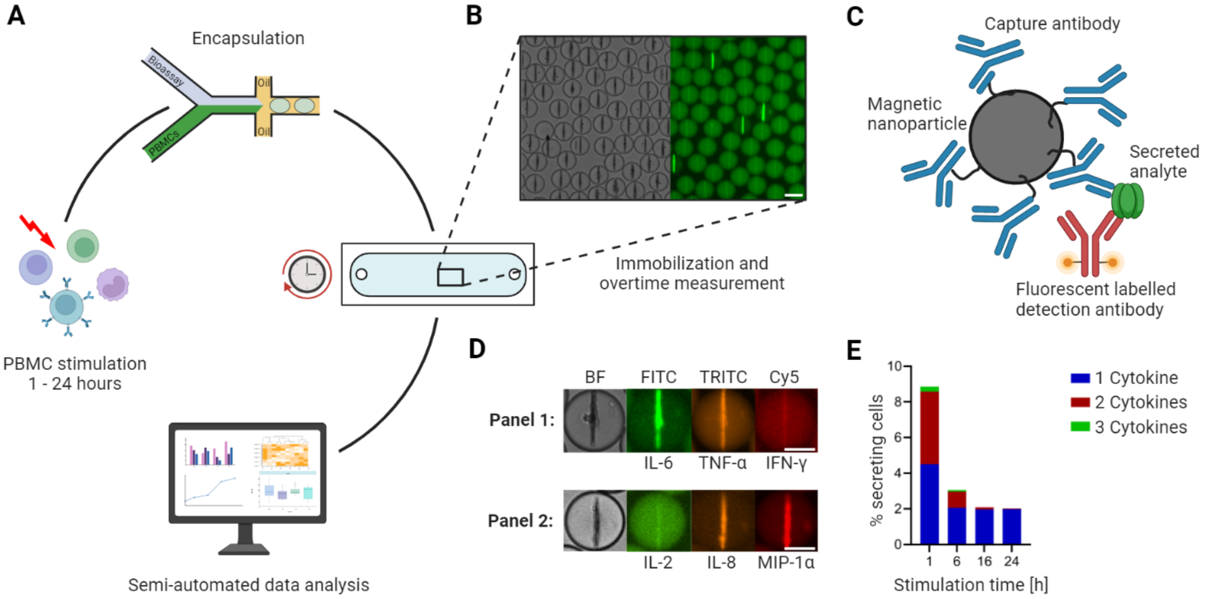


**Figure 1. Workflow of the microfluidic cytokine secretion measurements.** A. The experimental protocol involved the stimulation of PBMCs in bulk, followed by their encapsulation with assay reagents in 65 picolitre water/oil emulsions (called droplets). Subsequently, the droplets containing cells were immobilized in an observation chamber and imaged over 4 hours every 30 minutes, followed by a semi-automated data analysis pipeline. B. Micrographs of an array of droplets immobilized in the observation chamber. The insert shows the droplets in brightfield and one fluorescence channel. Droplets with lines indicate the presence of a cytokine secreting cell in this container. Scale bar: 50 µm. C. Assay principle to measure cytokine secretion. Assay reagents consist of 300 nm in diameter paramagnetic nanoparticles and fluorescently labeled detection antibodies. The nanoparticles are functionalized with capture antibodies specific to the cytokines of interest. When secreted, the cytokine binds to the capture antibody with subsequent relocation of one particular fluorescently labeled detection antibody. The application of a magnetic field aligns the nanoparticles into an elongated aggregate, making it possible to measure fluorescence relocation for every channel and to measure fluorescence relocation onto the nanoparticles (as seen in B). D. Nanoparticles functionalized against different cytokines allow multiplexing for up to three cytokines. The images shown represent exemplary cells secreting IL-6$^{+}$/TNF-α$^{+}$ (in a panel measuring IL-6/TNF-α/IFN-γ) and IL-8$^{+}$/MIP-1α$^{+}$ (IL-2/IL-8/MIP-1α). Scale bars: 25 µm. E. Response of LPS-stimulated PBMCs for various stimulation times, namely 1, 6, 16 and 24 hours. The resulting percentage of secreting cells was binned for polyfunctionality, i.e., cells secreting one (blue), two (red) or all three (green) measured cytokines (panel IL-6/TNF-α/IL-1β). Panels A and C created with BioRender.com.

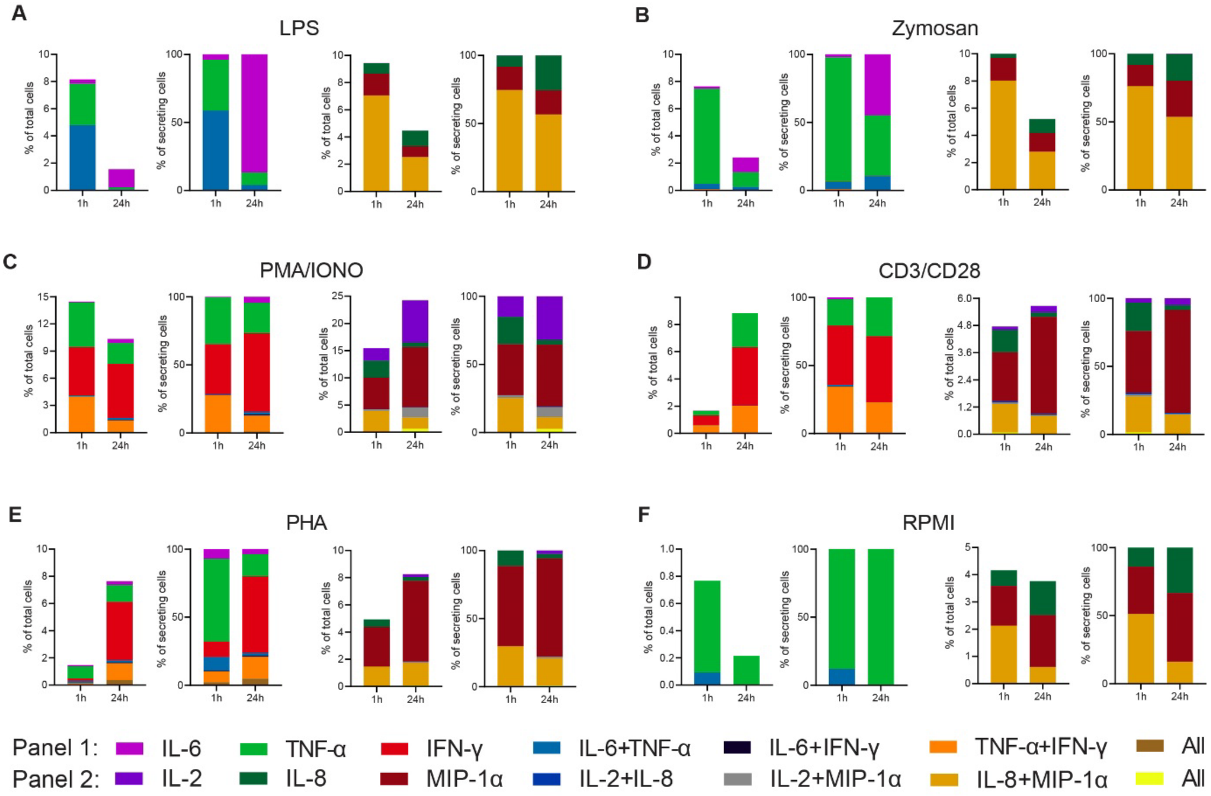


**Figure 2. Cytokine polyfunctionality of stimulated PBMCs changes with prolonged stimulation.** PBMCs were stimulated with either 1 µg/ml LPS (A), 100 µg/ml zymosan (B), 50 ng/ml PMA + 1 µg/ml ionomycin (C), 5 µg/ml anti-CD3 (OKT3)/anti-CD28 (CD28.2, D), 10 µg/ml PHA-L (E) or media alone (F) for 1 and 24 hours and subsequently measured over 4 hours. Endpoint percentage CSCs were categorized into the following bins depending on the detected cytokines per droplet and the detection panel used IL-6 (violet), TNF-α (green), IFN-γ (red), IL-6$^+$/TNF-α$^+$ (blue), IL-6$^+$/IFN-γ$^+$ (black), TNF-α$^+$/IFN-γ$^+$ (orange), all three cytokines (brown) and IL-2 (dark violet), IL-8 (dark green), MIP-1α (dark red), IL-2$^+$/IL-8$^+$ (dark blue), IL-2$^+$/MIP-1α$^+$ (grey), IL-8$^+$/MIP-1α$^+$, or all three cytokines (yellow). The number of secreting cells in each bin is displayed as frequency of total measured cells (left panels) and frequencies of all CSCs per measurement (right panels). $n_{biological\ replicates}=1$.

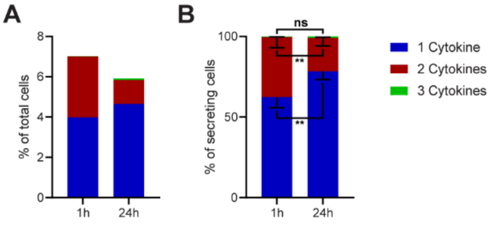


**Figure 3. Polyfunctionality of all measured cytokines and stimulants (summary of Figure 2).** All measured cytokines and stimulants were combined and binned into cells secreting one, two or three cytokines simultaneously. A. The average secreting cells in each bin over all the measurements in relation to all measured cells for 1- and 24-hour stimulations (see methods for details). B. Average of the normalized secreting cells in each bin in relation to all secreting cells. Differences between 1- and 24-hour stimulations were assessed with paired two-sided t-tests with 95% confidence. Data depicted as mean ± SEM. n=12.

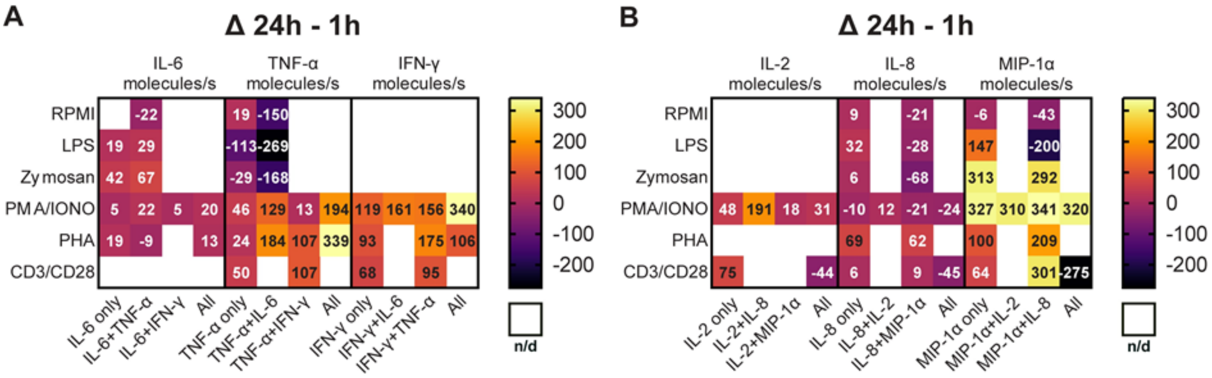


**Figure 4. Secretion rates of polyfunctional cells vary for different cytokines and stimulation times.** Normalized average secretion rates in molecules/s (1-hour subtracted from 24-hour measurement) over the measurement time (4 hours) for PBMCs stimulated with LPS, zymosan, PMA/ionomycin, anti-CD3/anti-CD28, PHA-L or media only for panel 1 (A) and panel 2 (B). Average secretion rates were extracted according to the different bins described in Figure 2. n/d = not detected.

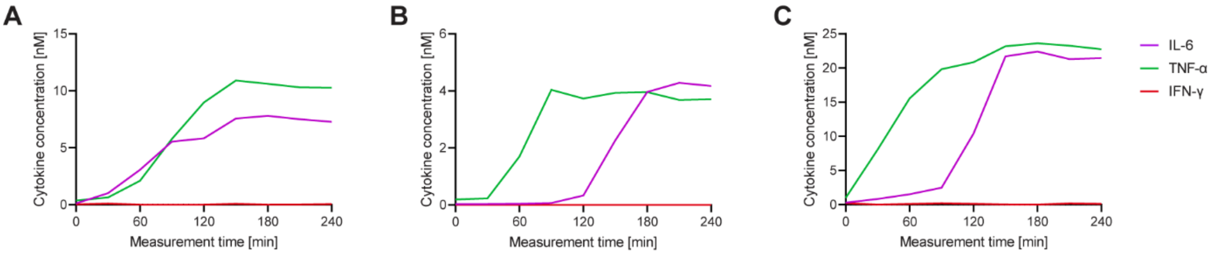


**Figure 5. Each cell displays a different secretion behavior after LPS stimulation for IL-6⁺/TNF-α⁺ polyfunctional cells.** Three exemplary cells secreting IL-6⁺/TNF-α⁺ after a 1-hour stimulation with 1 µg/ml LPS and subsequent single cell measurement for 4 hours (A-C). Measured cytokines included IL-6 (purple), TNF-α (green) and IFN-γ (red). Secretion dynamics are visualized by plotting measured in-droplet concentration against measurement time.

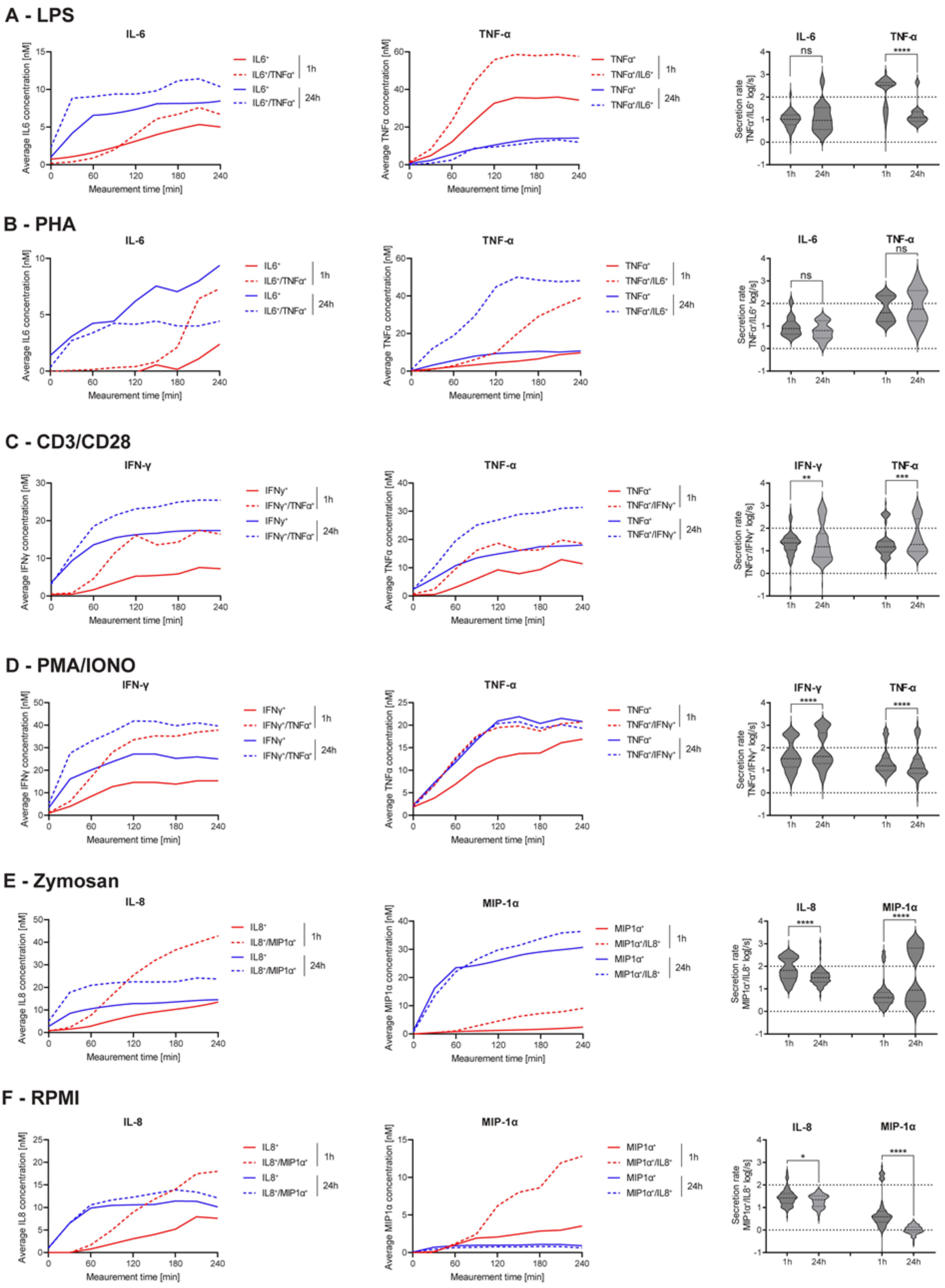


**Figure 6. Secretion dynamics change for polyfunctional cytokine secreting populations**. Secretion behavior changes for co-secreting cells with prolonged incubation times. IL-6$^{+}$/TNF-α$^{+}$ secreting cells in response to 1- and 24-hour stimulations with LPS (A) and PHA (B). TNF-α$^{+}$/IFN-γ$^{+}$ secreting cells in response to 1- and 24-hour stimulations with anti-CD3/anti-CD28 (C) and PMA/ionomycin (D). IL-8$^{+}$/MIP-1α$^{+}$ secreting cells in response to 1- and 24-hour stimulations with zymosan (E) and for non-stimulated cells (F). 1$^{st}$ and 2$^{nd}$ panels show the

average in-droplet concentrations of the respective populations over the measurement time (4 hours), compared to the single-cytokine secretion population. 3rd panels show the average secretion rate (log) distributions of individual IL-6$^+$/TNF-α$^+$ secreting cells over the whole measurement time for 1- and 24-hour stimulations and for both measured cytokines. Statistical differences in secretion rate distributions were assessed using two-sided, unpaired, nonparametric Kolmogorov-Smirnov t-tests with 95% confidence and $p < 0.05$. Data with less than ten detected secreting cells is not depicted. Vertical dotted lines show secretion rates of 1 and 100 molecules per second, respectively. $n_{\text{biological replicates}}=1$.

### Supplementary figures

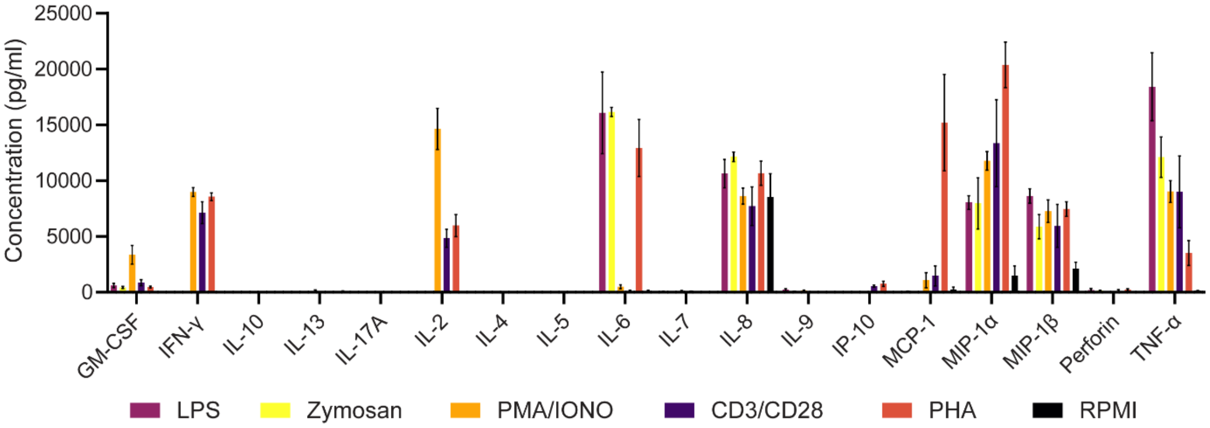


**Figure S1. Cytokine supernatant concentrations after 24 hours in response to various stimulants.** Average cytokine concentrations in the supernatant (mean ± SEM) after stimulation. Measured were the supernatant concentrations of GM-CSF, IFN-γ, IL-10, IL-13, IL-17A, IL-2, IL-4, IL-5, IL-6, IL-7 IL-8, IL-9, IP-10, MCP-1, MIP-1α, MIP-1β, Perforin and TNF-α after 24-hour stimulations with 1 µg/ml LPS (purple), 100 µg/ml zymosan (yellow), 50 ng/ml PMA + 1 µg/ml Ionomycin (orange), 5 µg/ml anti-CD3 (OKT3)/anti-CD28 (CD28.2, violet), 10 µg/ml PHA-L (light red) or media alone (black). N = 3.

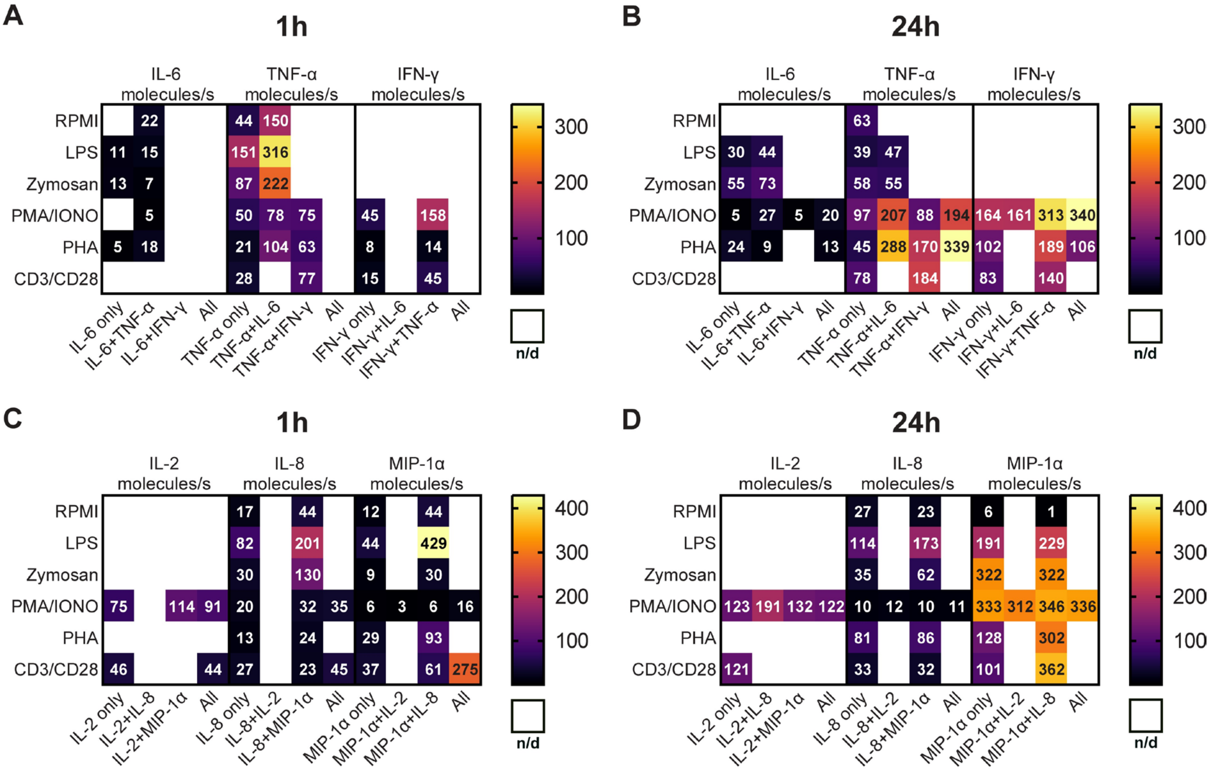


**Figure S2. Absolute secretion rates of polyfunctional cells vary for different cytokines and stimulation times (refers to Figure 4).** Average secretion rates in molecules/s over the measurement time (4 hours) for PBMCs stimulated with LPS, zymosan, PMA/ionomycin, anti-CD3/anti-CD28, PHA-L or media only for panel 1 (A, B) and panel 2 (C, D). Cell were stimulated for 1 and 24 hours. n/d = not detected.
